# Hydrogenotrophic methanogenesis in archaeal phylum Verstraetearchaeota reveals the shared ancestry of all methanogens

**DOI:** 10.1101/391417

**Authors:** Bojk A. Berghuis, Feiqiao Brian Yu, Frederik Schulz, Paul C. Blainey, Tanja Woyke, Stephen R. Quake

## Abstract

Methanogenic archaea are major contributors to the global carbon cycle and were long thought to belong exclusively to the euryarchaeotal phylum. Discovery of the methanogenesis gene cluster methyl-coenzyme M reductase (Mcr) in the Bathyarchaeota and thereafter the Verstraetearchaeota led to a paradigm shift, pushing back the evolutionary origin of methanogenesis to pre-date that of the Euryarchaeota. The methylotrophic methanogenesis found in the non-Euryarchaota distinguished itself from the predominantly hydrogenotrophic methanogens found in euryarchaeal orders as the former do not couple methanogenesis to carbon fixation through the reductive acetyl-coenzyme A (Wood-Ljungdahl) pathway, which was interpreted as evidence for independent evolution of the two methanogenesis pathways. Here, we report the discovery of a complete and divergent hydrogenotrophic methanogenesis pathway in a novel, thermophilic order of the Verstraetearchaeota which we have named *Candidatus Methanohydrogenales*, as well as the presence of the Wood-Ljungdahl pathway in the crenarchaeal order *Desulfurococcales*. Our findings support the ancient origin of hydrogenotrophic methanogenesis, suggest that methylotrophic methanogenesis might be a later adaptation of specific orders, and provide insight into how transition from hydrogenotrophic to methylotrophic methanogenesis might occur.

## Introduction

All known methanogenic organisms belong exclusively to the archaeal domain of life, and are typically found in the oxygen-depleted environments of soils, sediments, and the intestinal tract of humans and animals (Thauer, 2008; Hedderich, 2010). With an estimated combined annual production of 500 million tons of the greenhouse gas methane, methanogenic archaea are key contributors to the global carbon cycle and play an important role in climate change (Reeburgh, 2007; IPCC, 2014). Until recently, all known methanogens belonged to the Euryarchaeota and were categorized into two classes (Class I and Class II). The hypothesis that methane metabolism originated early in the evolution of the Euryarchaeota (Gibraldo, 2006) has since been challenged following the recent discovery of putative methane metabolism in the archaeal phyla Bathyarchaeota (formerly Miscellaneous Crenarchaeota Group) (Evans, 2015; Borrel, 2016) and Verstraetearchaeota (Vanwonterghem, 2016).

Three major pathways of methanogenesis are known (Ferry, 2007; Conrad, 2009): hydrogenotrophic, methylotrophic and acetoclastic (**Fig. 1A**). The only enzyme that is present in all types of methanogenesis is methyl-coenzyme M reductase (Mcr), a Ni-corrinoid protein catalyzing the last step of methyl group reduction to methane (Hedderich, 2013; Liu, 2008; Thauer, 2008). Hydrogenotrophic methanogenesis is the most widespread (Thauer, 2008) and has been suggested to represent the ancestral form of methane production (Bapteste, 2005). Class I methanogens (*Methanopyrales*, *Methanococcales*, and *Methanobacteriales*) as well as most of Class II methanogens (*Methanomicrobiales, Methanocellales and Methanosarcinales*, with the exception of the *Methanomassiliicoccales*) are hydrogenotrophs. They reduce CO_2_ to CH_4_ in six steps via the reductive acetyl-coenzyme A (acetyl-CoA) or Wood-Ljungdahl pathway (WLP) - one of the most important processes for energy generation and carbon fixation (Berg, 2010) - by using H_2_ or sometimes formate as an electron donor (Liu, 2008; Thauer, 2008). Conservation of energy takes place by coupling of the WLP to methanogenesis, with the methyl-tetrahydromethanopterin (methyl-H4MPT):coenzyme M methyltransferase complex (Mtr) being the key enzyme. Mtr uses the free energy of methyl transfer to establish a Na^2+^-motive force across the membrane (Schlegel, 2013) and low-potential reducing equivalents for CO_2_ reduction are provided by electron bifurcation at the cytoplasmic heterodisulfide reductase complex (HdrABC) (Kaster, 2011; Costa, 2010).

**Fig. 1.**
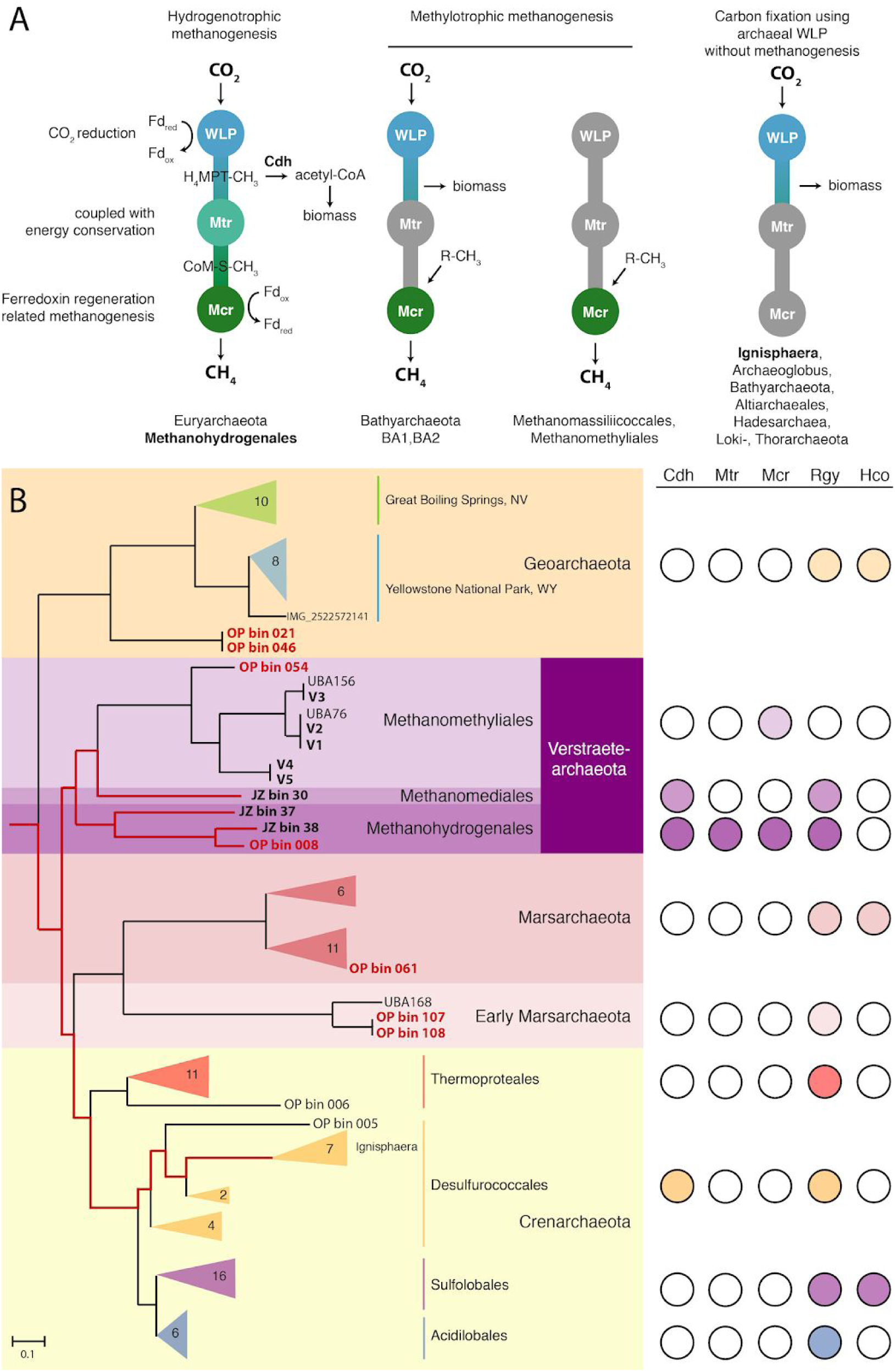
The Wood-Ljungdahl pathway coupled to methanogenesis in *Methanohydrogenales*. (**A**) Different configurations for the associated or independent functioning of the archaeal version of the Wood-Ljungdahl pathway (WLP) and methanogenesis. Missing enzymatic complexes or pathways are shaded in gray. From left to right: CO_2_-reducing methanogenesis as present in *Methanohydrogenales* as well as Class I and Class II methanogens without cytochromes; Methanogenesis by reduction of methyl-compounds using H_2_ as inferred in Bathyarchaeota BA1, and potential link with the WL pathway in absence of Mtr; Methanogenesis by reduction of methyl-compounds using H_2_ as present in *Methanomassiliicoccales* and *Methanomethyliales*; Carbon fixation using the archaeal WL pathway in absence of methanogenesis, and proposal of a mechanism to generate low potential ferredoxin during sulphate reduction in the case of Archaeoglobales. (**B**) Genome-based phylogeny (left) of the TACK superphylum genomes found in the Obsidian Pool (OP) dataset. The tree was inferred using a concatenated set of 56 marker genes, using the DPANN superphylum as the outgroup. OP genomes found in the Geo-, Verstraete- and Marsarchaeota phyla are shown in red alongside members or clades containing publicly available genomes; the Crenarchaeota contain only OP genomes. Ancestry of the WLP in the TACK superphylum (red branches) inferred through clades that retained the complete set of WLP homologues. Scale bar indicates substitutions per site. (right) The presence (filled) or absence (empty circle) of key genes in the TACK clades. Mcr: methyl-coenzyme M reductase; Mtr: tetrahydromethanopterin S-methyltransferase subunits A-H; Cdh: CO dehydrogenase/acetyl-coenzyme A synthase; Rgy: reverse gyrase; Hco: heme copper-oxidase subunits I+II.

Bathyarchaeota, Verstraetearchaeota and *Methanomassiliicoccales* are - together with some *Methanobacteriales* and *Methanosarcinales* - methylotrophic methanogens. Here methylated C_1_ compounds, including methanol, methylamines and methylsulfides, are first activated by specific methyltransferases (Lang, 2015). Next, one of four methyl groups is oxidized through the same reactions as in the hydrogenotrophic pathway occurring in reverse, and the remaining three groups are reduced to methane by Mcr. As the non-euryarchaeal methanogens have been found to be exclusively methylotrophic, it was hypothesized that methylotrophic methanogenesis is an independently evolved ancient pathway (Borrel, 2016; Williams et al. 2017).

Alongside the discovery of the key methanogenesis gene cluster Mcr outside the euryarchaeal phylum, the key WLP gene cluster, CO dehydrogenase/acetyl-CoA synthase (Cdh) and associated genes have been found to occur independently of the methanogenesis pathway (Adam et al. 2018). While first thought to uniquely exist in the non-methanogenic euryarchaeal order *Archaeoglobales* as a remnant of its ancestral methane-cycling lifestyle (Vorholt et al. 1995; Bapteste et al. 2005), the presence of the archaeal WLP in the absence of Mcr and Mtr complexes has now been reported from an increasing number of archaeal lineages within the Bathyarchaeota (Sara Lazar et al. 2015; He et al. 2016), the Altiarchaeales (Probst et al. 2014), the Hadesarchaea/MSBL-1 (Baker et al. 2016; Mwirichia et al. 2016), the Lokiarchaeota (Sousa et al. 2016) and the Thorarchaeota (Seitz et al. 2016), with likely more awaiting discovery. While mechanisms to generate low potential reduced ferredoxin for CO_2_-reduction by the archaeal WL pathway in the absence of methanogenesis remain to be elucidated (Borrel 2016), the hypothesis that WLP-coupled methanogenesis is unique to the Euryarchaeota seemed to hold true (Sorokin et al. 2017).

To further explore the evolutionary diversity of methanogenic organisms, we collected and studied 5 samples from the Obsidian Pool (OP) hot spring in Yellowstone National Park (**Fig. S1, Table S1**). We were able to assemble 111 metagenome-assembled genomes (MAGs) through mini-metagenomic (Yu et al., 2017) and metagenomic analysis of these samples. These genomes are predominantly archaeal (98 out of 111, named OP bin 000 - 110) and most of the MAGs discovered in these samples were affiliated with the TACK superphylum (**Fig. 1B**). A number of MAGs belonging to the strictly anaerobic crenarchaeal family *Ignisphaera* contain all WLP genes, suggesting that some Crenarchaeota are in fact capable of carbon fixation. One of the Verstraetearchaeota (OP bin 008) forms a deeply-branching clade together with two recently published genomes from a hot spring sediment sample in Tengchong, Yunnan, China (JZ bins 37 and 38). Surprisingly, this clade contains all genes coding for the full pathway of hydrogenotrophic methanogenesis coupled to carbon fixation through the WLP. This finding places the origin of hydrogenotrophic methanogenesis outside the euryarchaeotal phylum, suggests that hydrogenotrophic and methylotrophic methanogenesis are more closely linked than once thought, and sheds light on how a transition from hydrogenotrophic to methylotrophic methanogenesis might have occurred.

### Expansion of the TACK superphylum by novel archaeal MAGs

A genome tree comprising the 98 Obsidian Pool (OP) archaeal genomes from our data and 1427 publicly available archaeal genomes was inferred using a concatenated set of 56 or 122 archaeal-specific marker genes (**Figs. 1B, S1**). The majority (49) of novel genomes fall within the TACK superphylum, of which 42 are Crenarchaeota. Our analysis extends the archaeal tree of life by adding novel lineages to the Bathy-, Geo-, Mars- and Verstraetearchaeota (**Fig. 1B; Table 1**). A KEGG orthology (Kaneshisa and Goto, 2000), COG (Tatusov et al., 2000), and pfam (Finn et al., 2014) based comparative analysis was performed on the OP archaeal bins and other members of clades containing novel lineages (**Figs. 1B, S2-4**). OP bin 061 groups within the newly reported Marsarchaeota (**Fig. 1B**, Jay et al., 2018). This placement is supported by functional analysis of the respective MAGs showing high similarity to traits typically found in the Marsarchaeota (**Fig. S2**). In contrast, OP MAGs 107 and 108, as well as UBA168, which branch basally to the Marsarchaeota, lack both heme-copper cytochrome oxidases (Hco) observed in other Marsarchaeota, suggesting anaerobic metabolism in this subclade. OP bin 46, an early branching Geoarchaeotum, contains both Hco subunits whereas all other (20) clade members contain only subunit 1, suggesting aerobic metabolism. OP bin 54 groups together with the Verstraetearchaeota. Though the genome is incomplete (**Table 1**), it contains subunits A,B,G,D of the Mcr gene cluster characteristic of this methylotrophic methanogenic phylum. OP bin 008 groups monophyletically with JZ bins 37 and 38 as an early branching clade of the Verstraetearchaota (**Fig. 1B**). While OP bin 008 lacks the 16S rRNA gene and JZ bin 38 contains an incomplete (502 bp) 16S rRNA sequence, JZ bin 37 full length 16S rRNA sequences show 84% identity to Verstraetearchaeota V1-3, 85% identity to V4 and 86% JZ bin 30 (**Fig. S6**). Partial 16S rRNA gene alignment of JZ bin 37 and 38 show 90% identity, and JZ bin 38 shows 83% and 81% identity to JZ bin 30 and V1-3, respectively. Based on these observations, we propose two new thermophilic orders in the Verstraetearchaeota, grouping JZ bins 38 and 37, as well as OP bin 008 as representatives of the first (*Candidatus Methanohydrogenales*), and JZ bin 30 as the first representative of the second (*Candidatus Methanomediales*).

**Table 1.** Summary statistics of novel archaeal phylogeny in the OP dataset.

| Genomic bin | Phylogeny | Assembly size (bp) | Contig N50 (bp) | # Contigs | GC-content | Completeness (%)* | Contamination (%)* |
| --- | --- | --- | --- | --- | --- | --- | --- |
| OP bin 008 | Verstraetearchaeota | 1,665,326 | 28,180 | 82 | 0.55 | 97.06 | 0.74 |
| OP bin 010 | Bathyarchaeota | 1,422,446 | 19,095 | 88 | 0.42 | 84.11 | 0.93 |
| OP bin 021 | Geoarchaeota | 1,147,812 | 12,080 | 106 | 0.32 | 55.23 | 0 |
| OP bin 042 | Bathyarchaeota | 1,286,079 | 16,585 | 96 | 0.42 | 87.2 | 1.87 |
| OP bin 046 | Geoarchaeota | 1,804,512 | 157,431 | 23 | 0.32 | 94.49 | 0.74 |
| OP bin 107 | Early Marsarch. | 84,454 | 6,214 | 13 | 0.55 | 21.99 | 0 |
| OP bin 054 | Verstraetearchaeota | 337,598 | 7,976 | 43 | 0.49 | 17.76 | 0 |
| OP bin 061 | Marsarchaeota | 2,310,778 | 51,573 | 84 | 0.44 | 96.32 | 4.41 |
| OP bin 108 | Early Marsarch. | 1,536,417 | 19,661 | 99 | 0.55 | 85.44 | 3.74 |

\* as estimated by CheckM (Parks et al., 2014)

### Hydrogenotrophic methanogenesis genes in Verstraetearchaeota, Wood-Ljungdahl pathway in Crenarchaeota

Comparative genomics of the two novel orders to the other verstraetearchaeotal order *Methanomethyliales* revealed that the former share 664 of the 838 Methanomethyliales homologues (of COG genes >30% identity), and have 342 additional unique gene homologues not found in the *Methanomethyliales*. The new orders, contrary to the other Verstraetearchaeota, have been sampled in hot springs and as such encode a reverse gyrase. Like the *Methanomethyliales*, they do not possess the Hco gene cluster, supporting an anaerobic metabolism. Further metabolic reconstruction of the *Methanohydrogenales* genomes revealed the presence of key genes associated with hydrogenotrophic methanogenesis via the archaeal Wood-Ljungdahl pathway (WLP, **Figs. 2A, 3A**). The methanogenic genes include methyl-coenzyme M reductase complex (Mcr; McrABGCD), all subunits of tetrahydromethanopterin S-methyltransferase (Mtr; MtrABCDEGH), and heterodisulfide reductase (Hdr; HdrABCD), and are found exclusively in the *Methanohydrogenales*. The WLP genes include CO dehydrogenase/acetyl-CoA synthase (Cdh; CdhABGDE), formylmethanofuran dehydrogenase (Fwd; FwdABCDE) and coenzyme F420-reducing hydrogenase (Frh; FrhABDG) and are present in both the *Methanohydrogenales* as the *Methanomediales*. In contrast to the methylotrophic methanogenic *Methanomethyliales* (Vanwonterghem *et al*. 2017), the *Methanohydrogenales* have retained all enzymes in the reductive acetyl-coenzyme A pathway as well as all subunits of Mtr and HdrABC (**Fig. 2A**). This suggests that the *Methanohydrogenales* are able to perform hydrogenotrophic methanogenesis, a metabolism that has until present eluded discovery outside the Euryarchaeota.

**Figure 2.**
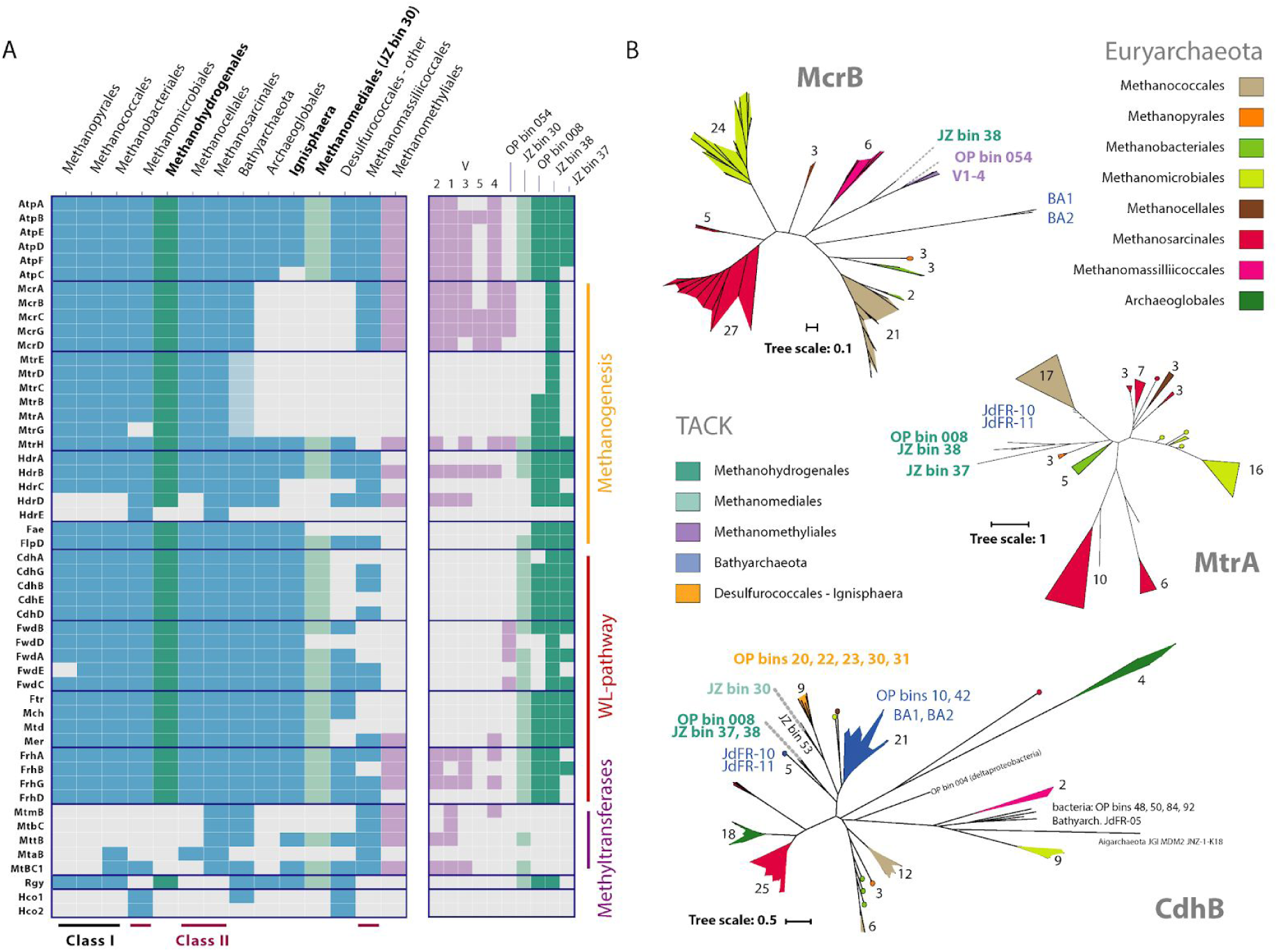
Key WLP and methanogenesis genes in Obsidian Pool genomes and clades, compared to other methanogens. (A) (left) Presence of genes in the clades of the 7 euryarchaeal methanogen groups, *Methanomethyliales* (purple), Bathyarchaeota, *Archaeoglobales*, *Ignisphaera*, other *Desulfurococcales*, *Methanomediales* (JZ bin 30, light green) and *Methanohydrogenales* (green), ordered by similarity (see **Table S2** for gene details). The full Mtr gene cluster was observed in two Bathyarchaeota (light blue), but the presence of the full hydrogenotrophic methanogenesis pathway in a single genome has not been observed (**Fig. S4**). (right) Single-genome presence of genes in the *Methanomethyliales* (purple), *Methanomediales* (light green), and *Methanohydrogenales* (green). See **Fig. S3** for additional genes. (B) Gene trees showing the phylogeny of CdhB, McrB and MtrA genes found in the OP MAGs relative to the 7 existing methanogenic euryarchaeal orders, the Archaeoblobales, Desulfurococcales and Bathyarchaeota (Additional subunit trees shown in **Figs S8-S13**). Numbers indicate the number of genomes present in a branch.

The genes for the reductive acetyl-CoA pathway are also observed in the *Ignisphaera*, a genus in the crenarchaeal family *Desulfurococcaceae* (**Figs. 1B, 2A**). The *Desulfurococcales*, a predominantly anaerobic hyperthermophilic order, are known to have the capacity for autotrophic carbon fixation through the dicarboxylate-hydroxybutyrate cycle. OP bins 20, 22, 23, 30 and 31 all contain Cdh, Fwd, Ftr, and Fhr (**Figs 1B, S4**). This suggests that they are capable of autotrophic carbon fixation through the WLP in a manner similar to the *Archaeoglobales* and other lineages. Contrary to most members of the *Desulfurococcales*, the Ignisphaera lack several enzymes associated with the dicarboxylate-hydroxybutyrate cycle (**Fig. S2**) such as pyruvate phosphate dikinase, Succinyl-CoA synthethase, Malate/lactate deydrogenase, as well as Fumarate reductase and dehydrogenase. Key enzyme 4-hydroxybutyryl-CoA dehydratase was also not found. These enzymes are observed in the other families within this order.

The *Methanohydrogenales* do not contain genes for methanogenesis from methylated compounds (**Fig. 2A**), unlike several euryarchaeal orders as well as the *Bathyarchaeota, Archaeoglobales* and *Desulfurococcales*. JZ bin 30, the only representative of the *Methanomediales*, does contain trimethylamine methyltransferase MttB, though it lacks Mtr and Mcr. The genetically encoded amino acid pyrrolysine was only observed in the methyltransferases (MtmB, MtbC and MttB, **Fig. S15**) of the *Methanosarcinales* and *Methanomassilliicoccales*, as previously reported (Borrel et al., 2014). The *Methanohydrogenales*, together with the *Ignisphaera*, many Bathyarchaeota and Euryarchaeota from all classes contain a formaldehyde activating enzyme necessary for methanogenesis (Fae), while the *Methanomethyliales* lack this gene. As in the *Methanomethyliales*, homologues of the membrane-bound NADH-ubiquinone oxidoreductase (Nuo, **Fig. S3**) are present and capable of re-oxidizing reduced ferredoxin as occurs in the Methanosarcinales (Welte and Deppemeyer, 2014), with concomitant translocation of protons or sodium ions across the membrane (**Fig 3A**). The use of ferredoxin as an electron donor further supported by the fact that the subunits required for the binding and oxidation of NADH (NuoEFG) are missing (**Fig. S3**). Ni,Fe hydrogenase subunits belonging to energy-conserving hydrogenase B (EhbMN, **Fig. S3**) are not present in the *Methanohydrogenales* and *Methanomediales*, contrary to the *Methanomethyliales*. However, the large and small subunits of Ni,Fe hydrogenase III (Nfh, **Fig. S3**) are present, suggesting that this complex might facilitate the generation of low potential electrons required for autotrophic carbon fixation (Major et al., 2010). This in turn can drive ATP synthesis via the archaeal type ATP synthase (AtpA-F). *Methanohydrogenales* lack several transporters of complex molecules that are found in the *Methanomethyliales*. *Methanomethyliales* contain genes for importing complex sugars, lipopolysaccharides and oligopeptides (**Figs. 3A, S5**). The ability of the *Methanomethyliales* to carry out complex fermentation, while *Methanohydrogenales* does not is likely a result of substrate diversification due to the rich environment of (active sludge) of the latter compared to the former.

**Fig. 3.**
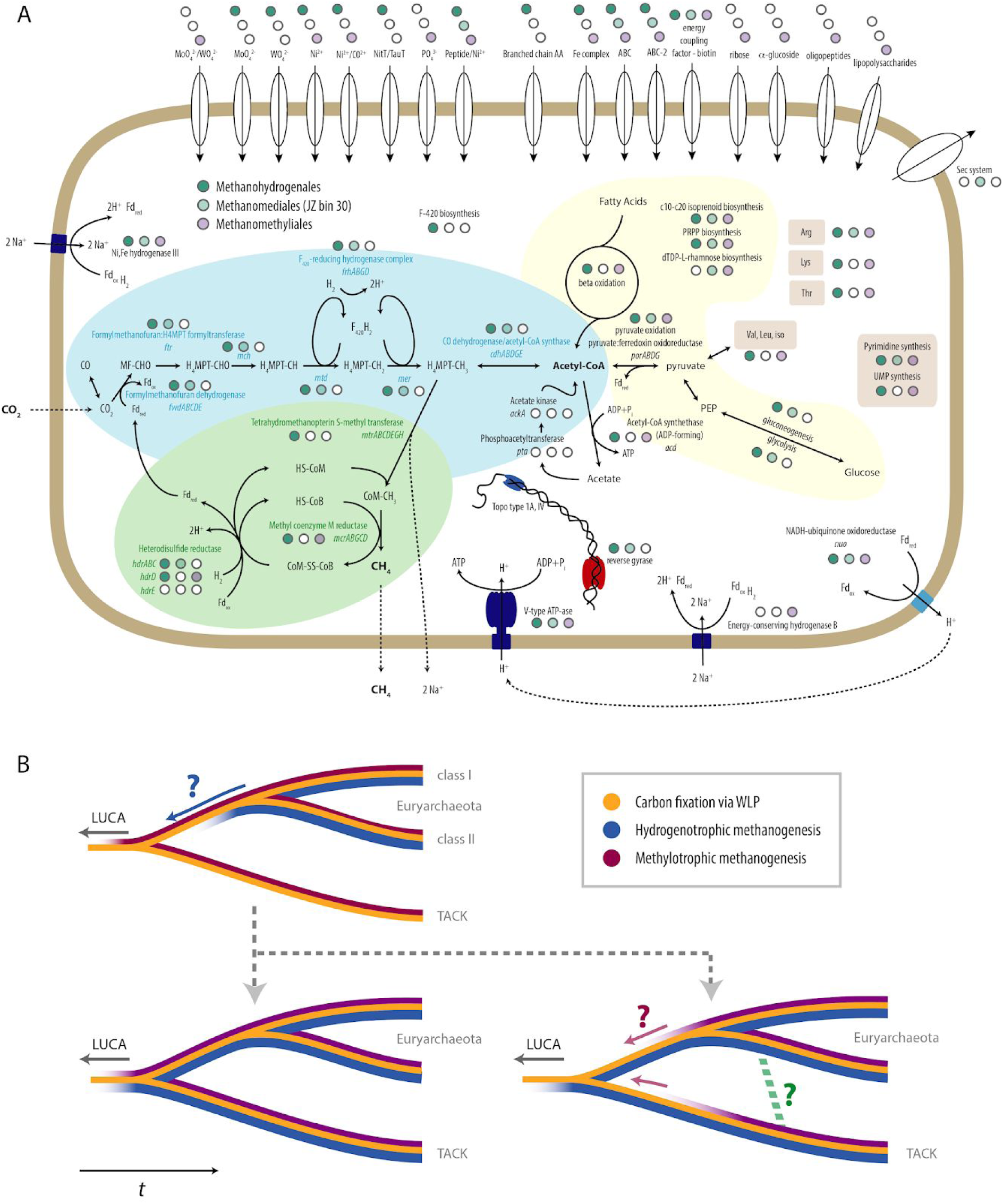
Metabolism of hydrogenotrophic methanogen Methanohydrogenales and the evolution of methanogenesis in archaea. (A) Proposed methanogenesis-related pathways and other metabolism of the three verstraetearchaeotal orders. Presence (colored circle) or absence (white circle) of a gene (cluster) is shown separately for the *Mehtanohydrogenales* (green), *Methanomediales* (light green), and the *Methanomethyliales* (purple). The full pathway for autotrophic carbon fixation via the archaeal-type Wood-Ljungdahl pathway (WLP, blue background), ferredoxin regeneration-related methanogenesis (green background), as well as metabolisms of carbohydrates and lipids (yellow background) or nucleotides and amino acids (brown) are shown. (B) (top) Hydrogenotrophic (or CO_2_-reducing) methanogenesis (blue) was previously exclusively observed in the Euryarchaeota, while methylotrophic methanogenesis (purple) was also found in the TACK superphylum. (bottom left) The discovery of the full hydrogenotrophic pathway in the Verstraetearchaeota - a member of the TACK superphylum - suggests that this pathway was present in the last common ancestor of the Verstraetearchaeota and Euryarchaeota. This observation also supports the hypothesis that methylotrophic methanogenesis is a later adaptation (including possible horizontal transfer of methyltransferases; green dashed line) of the more basal CO_2_-reducing form, and further strengthens the notion that the last universal common ancestor (LUCA) was a CO_2_-reducing methanogen.

### Evolutionary history of hydrogenotrophic methanogenesis in Verstraetearchaeota

The full hydrogenotrophic methanogenesis pathway has not been previously identified in microorganisms outside Euryarchaeota. To further explore the distribution and diversity, phylogenetic trees of Mcr, Mtr and Cdh (or subunits thereof) were calculated (**Figs. S8-S14**). Phylogenetic analysis of MtrA, B and G, the three subunits in addition to MtrH that are present in bin 008 and JZ bin 38, resulted in a robust grouping of these two genomes in a monophyletic clade (**Figs. 2B, S8-10**) together with the two Bathyarchaeota genomes containing Mtr (JdFR-10, JdFR-11, **Fig. 2B, S4**) branching off at the root of the euryarchaeota. This non-euryarchaeal clade is closely affiliated with the *Methanobacteriales* and (in the case of MtrA and B) the *Methanopyrales*. The trees of McrA and McrB show that OP bin 54 branches off closely to the Verstraetearchaeota (V1-5, **Figs. 2B, S11**). JZ bin 38, the only genome in *Methanohydrogenales* that possesses the Mcr gene cluster, is present on the same branch as the *Methanomethyliales*/other Verstraetearchaeota, suggesting a similar evolutionary trajectory of this gene cluster. JZ bin 38 branches in basal position to OP bin 54 for McrB, and for McrA OP bin 54 branches basally to JZ bin 38, together with the McrA genes of Bathyarchaeota BA1 and BA2. The CdhB, D, and G gene trees show more diversity in the TACK superphylum and the Bathyarchaeota as a result of the more common presence of the Cdh gene cluster throughout these clades (**Figs. 2B, S12,13**). For McrB, the three *Methanohydrogenales* are closely related to JdFR-10 and JdFR-11, the two Bathyarchaeota with the full Mtr gene cluster (**Fig. 2B**). This clade forms the first branch of a group containing all Bathyarchaeota, the *Ignisphaera* and JZ bin 30. *Methanohydrogenales* have more divergent CdhD, yet form a single clade. JZ bin 30 branches ancestral to a clade containing the Ignisphaera in CdhB, D and G, showing divergent evolution of the acetyl-CoA synthase gene cluster in the Crenarchaeota as well as branches leading toward the Verstraetearchaeota. CdhG shows that JZ bin 37 and Bathyarchaeota JdFR-10 and 11 diverge from the position of the other *Methanohydrogenales* as an early branch of the TACK superphylum and Bathyarchaeota, showing a closer affiliation to *Methanopyrales*.

The presence of subunit H of the Mtr gene cluster correlates with the presence of the gene clusters associated with autotrophic carbon fixation through the Wood-Ljungdahl pathway. As such, MtrH shows a distinctly different evolutionary pattern than subunits A-G (**Fig. S10**). It has been suggested that it plays a role in methylamine:coenzyme M methyl transfer activity in the *Methanomethyliales* (Vanwonterghem et al. 2016), yet it is unclear what its role is in non-methanogens where it is observed, such as the *Archaeoglobales*, *Ignisphaera* and many Bathyarchaeota. Other genes typically associated with hydrogenotrophic methanogenesis are found throughout members of the Bathyarchaeota (Mtr in JdFR-10 and JdFR-11; **Fig. S4, S7**), Archaeoglobales (Fae, HdrA-D, MtrH), Ignisphaera (Fae, HdrAB, MtrH, EhbCDHI; **Fig. 2A, S7**). In addition, OP bin 54, a member of the *Methanomethyliales*, contains an almost intact formylmethanofuran dehydrogenase complex (FwdABCD, **Fig. 2A**), a key enzyme complex in the WLP. It is plausible that the presence of these genes might constitute remnants of ancestral hydrogenotrophic methanogenesis, as the various lineages described here adopted to other modes of energy production and metabolism.

The discovery of two novel orders of hyperthermophilic Verstraetearchaeota sheds light on how the transition from autotrophic hydrogenotrophic methanogenesis to heterotrophic methylotrophic methanogenesis might have occurred. Decoupling of carbon fixation through the WLP and methane production might have been achieved through the acquisition of methyltransferases, rendering the Mtr complex redundant while maintaining the methanogenic mode of energy conservation. Signatures for this hypothesis can be found in the observation of two methyltransferases in the *Methanomediales* (JZ bin 30), the order more closely related to the *Methanomethyliales*, while the *Methanohydrogenales* lack methyltransferases (**Fig. 2A**). Signatures of a transition to other modes of non-methanogenic autotrophic growth can be observed within the Crenarchaeota, as we now observe the preservation of the WLP in the Ignisphaera, while other members of the hyperthermophilic Desulfurococcales and Thermoproteales are thought to be capable of autotrophic carbon fixation through the dicarboxylate-hydroxybutyrate cycle (Berg, 2010).

The congruent topology of the species and gene trees, as well as the presence of the entire collection of genes of the hydrogenotrophic methanogenesis pathway (**Fig. 3A**) suggest that these complexes have evolved as functional units and that hydrogenotrophic methanogenesis was present in the last common ancestor of the Verstraetearchaeota and Euryarchaeota (**Fig. 3B)**. In addition, the presence of the hydrogenotrophic pathway in the TACK superphylum makes it plausible that methylotrophic methanogenesis is the later adaptation. While the methylotropic pathway requires only a subset of the genes from the hydrogenotropic pathway, the hydrogenotropic pathway enables metabolism from a much simpler carbon source. This leaves us with genetic evidence for two possible evolutionary origins of methanogenesis (**Fig. 3B**): in the first scenario, hydrogenotrophic methanogenesis first evolved to support life in a nutrient poor environment that required using only CO_2_ as a source of carbon and then the simpler methanotropic pathways evolved through gene deletion as more complex nutrient environments became available, while in the second scenario methanotropic methanogenesis first evolved in a nutrient-rich environment where complex organic molecules were readily available and then the metabolic potential expanded through further gene addition as organisms eventually colonized more nutrient poor environments.

## Methods

### Environmental sample collection and storage

The environmental samples used in this study were collected from two separate hot springs in Yellowstone National Park under permit number YELL-2009-SCI-5788 (**Table S1**). The samples were collected from sediments of the Obsidian Pool in the mud volcano area and placed in 2 mL tubes without any filtering and soaked in 50% ethanol onsite (or Glycerol). After mixing with ethanol, samples were kept frozen until returning from Yellowstone to Stanford University, at which time tubes containing the samples were transferred to -80˚C for long term storage.

### (Mini-)metagenomic sequencing

#### Sample preparation and dilution for mini-metagenomic pipeline

Each sample from Yellowstone was thawed on ice. The tube was vortexed briefly to suspend cells but not large particles and debris. 1 mL of sample from the top of the tube was removed, placed in a new 1.5 mL tube, and spun down at 5000 x g for 10 min to pellet the cells. Supernatant was removed and cells were resuspended in 1% NaCl. After resuspension, cell concentration was quantified using a hemocytometer under bright field and phase microscopy (Leica DMI 6000). Each sample was then diluted in 1% NaCl or PBS to a final concentration of ~2.10^6^ cells/mL, corresponding to ~10 cells per chamber on the Fluidigm C1 microfluidic IFC (Integrated Fluidic Circuit).

#### Microfluidic genomic DNA amplification on Fluidigm C1 auto prep system

Because of the small amount of DNA associated with 5–10 cells, DNA contamination was a concern for MDA reactions. To reduce DNA contamination, we treated the C1 microfluidic chip, all tubes, and buffers under UV (Strategene) irradiation for 30 min following suggestions of Woyke *et al*. (Woyke et al., 2011). Reagents containing enzymes, DNA oligonucleotides, or dNTPs were not treated. After UV treatment, the C1 IFC was primed following standard protocol (https://www.fluidigm.com/binaries/content/documents/fluidigm/resources/c1-dna-seq-pr-100-7135/c1-dna-seq-pr-100-7135/fluidigm%3Afile). Priming the C1 IFC involved filling all microfluidic control channels with C1 Harvest Reagent, all capture sites with C1 Blocking Reagent, and the input multiplexer with C1 Preloading Reagent. These reagents were available in all C1 Single-Cell Reagent Kits. The diluted environmental sample was loaded onto the chip using a modified version of the loading protocol where washing was not performed, as the capture sites were too large for microbial cells. Hence, they acted essentially as chambers into which cells were randomly dispersed (**Fig. S1A**). Following cell loading, whole genome amplification via MDA was performed on-chip in 96 independent reactions. A lysozyme (Epicenter) digestion step was added before alkaline denaturation of DNA. After alkaline denaturation of DNA, neutralization and MDA were performed (Qiagen REPLI-g single cell kit). Concentrations of all reagents were adjusted to match the 384 well plate-based protocol developed by the single-cell group at DOE’s Joint Genome Institute but adapted for volumes of the Fluidigm C1 IFC (Rodrigue et al., 2009).

Lysozyme digest was performed at 37˚C for 30 min, alkaline denaturation for 10 min at 65˚C, and MDA for 2 hr and 45 min at 30˚C. The detailed custom C1 scripts for cell loading, DNA amplification, and associated protocols including reagent compositions are available through Fluidigm’s ScriptHub at the following link: https://www.fluidigm.com/c1openapp/scripthub/script/2017-02/mini-metagenomics-qiagen-repli-1487066131753-8

#### DNA quantification, library preparation and sequencing

Amplified genomic DNA from all sub-samples was harvested into a 96 well plate. The concentration of each sub-sample was quantified in independent wells using the 96 capillary version of the Fragment Analyzer from Advanced Analytical Technologies Inc. (AATI). Because of the size of recovered DNA, we used the high sensitivity large fragment analysis kit (AATI) and followed its standard protocol. The instrument effectively runs 96 independent gel electrophoresis assays, producing an electropherogram for each sub-sample. Using a DNA ladder with known concentrations as reference, the instrument’s software quantifies DNA concentration for each sub-sample by performing smear analysis between user specified ranges. For MDA amplified microbial genomic DNA harvested from Fluidigm’s C1 IFC, one large smear was often present between 1.5 kbp and 30 kbp. Following quantification, DNA from each sub-sample was diluted to 0.1–0.3 ng/mL, the input range of the Nextera XT library prep pipeline. Nextera XT V2 libraries (Illumina) were made with dual sequencing indices, pooled, and purified with 0.75 volumes of AMpure beads (Agencourt). Illumina Nextseq (Illumina) 2×150 bp sequencing runs were performed on each library pool.

#### Shotgun metagenomics

Bulk genomic DNA were extracted from Yellowstone National Park hot spring samples using Qiagen’s blood and tissue kit using the protocol for DNA extraction from gram positive bacteria. Nextera V2 libraries were constructed and sequenced on Illumina’s Nextseq platform. 24, 20, 49, 6.6 and 104 million reads were obtained from Obsidian Pool samples 1-5 respectively (**Table S1**), and trimmed using the same parameters as the mini-metagenomic sequencing reads. Finally, assembly is performed using Megahit (Li et al., 2015), with default options and kmer values of 21, 31, 41, 51, 61, 71 and 99, as well as MetaSPAdes (Nurk et al., 2017)

### Sequence assembly and annotation

A custom bioinformatic pipeline was used to generate combined biosample contigs (Yu et al., 2017). Sequencing reads were filtered with Trimmomatic (V0.30) in paired end mode with options ‘ILLUMINACLIP:adapters.fa:3:30:10:3:TRUE SLIDINGWINDOW:10:25 MAXINFO:120:0.3 LEADING:30 TRAILING:30 MINLEN:30’ to remove possible occurrences of Nextera indices and low quality bases (Bolger et al., 2014). Filtered reads from each sub-sample were clustered using DNACLUST, with k = 5 and a similarity threshold of 0.98, in order to remove reads from highly covered regions (Ghodsi et al., 2011). Then, assembly was performed using SPAdes (V3.5.0) with the sc and careful flags asserted (Bankevich et al., 2012). From all sub-sample assembly output, corrected reads were extracted and combined. The combined corrected reads from all sub-samples were assembled again via SPAdes (V3.5.0) with kmer values of 33,55,77,99. Finally contigs longer than 5 kbp were retained for downstream analyses.

The contigs were uploaded to JGI’s Integrated Microbial Genomes’ Expert Review online database (IMG/ER) for annotation (Huntemann et al., 2016). Briefly, structural annotations were performed to identify CRISPRs (pilercr), tRNA (tRNAscan), and rRNA (hmmsearch). Protein coding genes were identified with a set of four *ab initio* gene prediction tools: GeneMark, Prodigal, MetaGeneAnnotator, and FragGeneScan. Finally, functional annotation was achieved by associating protein coding genes with COGs, Pfams, KO terms, EC numbers. Phylogenetic lineage was assigned to each contig based on gene assignments.

### Binning, reassembly, pruning and quality assessment

Contig binning was performed by creating a 2D projection of the contig 5-mer space using the dimensionality reduction algorithm tSNE (Van der Maaten and Hinton, 2008), followed by unsupervised clustering of grouped contigs using HDBscan (McInnes et al., 2017). The redundancy of these bins, often containing a mix of contigs from the three different assemblers used (SPAdes for mini-metagenomic and MetaSPAdes or Megahit for the metagenomic contigs), was reduced by reassembling all the reads mapping onto the contig of a particular bin using MetaSPAdes. The resulting reassembled bin was then further pruned or split based on GC-content and coverage similarity. The bin quality was assessed using CheckM; only bins with contamination <5% and strain heterogeneity <1% were used for further analysis.

### Genome tree phylogeny

The 98 archaeal MAGs generated in this study were placed in phylogenetic context with 4472 publicly available archaeal genomes available in IMG/M (Chen et al., 2016, database accessed March 2018) and recently published Marsarchaeota (Jay et al., 2018) using two sets of conserved phylogenetic marker proteins: 56 universal proteins (Yu et al., 2017) or 122 archaeal-specific marker genes (http://gtdb.ecogenomic.org/). In brief, marker proteins were identified with hmmsearch (version 3.1b2, hmmer.org) using a specific hmm for each of the markers. For every protein, alignments were built with MAFFT (Katoh and Standley, 2013) (v7.294b) and subsequently trimmed with trimAl 1.4 (Capella-Gutierrez et al., 2009), removing sites at which more than 90 percent of taxa contained missing information. Genomes with less than 5 marker proteins and genomes which had more than 5 duplicated marker proteins were removed from the alignment. Based on an initial tree built with FastTree2 (REF, -spr 4 -mlacc 2 -slownni -lg) a subset of archeal MAGs and reference genomes affiliated with Geoarchaeota, Verstraetearchaeota, Marsarchaeota was selected. These genomes were then used to build the final phylogenetic tree with IQ-tree and the ultrafast bootstrap option (Nguyen et al., 2015, v1.5.5, REF, LG+F+I+G4 -alrt 1000 -bb 1000) on an alignment with 11134 informative sites for the concatenated alignment of the 56 universal marker proteins and 26962 informative sites for the concatenated alignment of the 122 archaeal marker proteins.

### Genome-wide and 16S rRNA average nucleotide identity

The 16S rRNA sequence average nucleotide identity (ANI) was calculated using the BLAST Global align Needleman-Wunsch algorithm (https://blast.ncbi.nlm.nih.gov/Blast.cgi?PAGE_TYPE=BlastSearch&PROG_DEF=blastn&BLAST_PROG_DEF=blastn&BLAST_SPEC=GlobalAln&LINK_LOC=BlastHomeLink). Genome-wide ANI was calculated using the ANI calculation tool of the IMG-ER web-based (meta)genome analysis toolset.

### Gene tree phylogeny

Extracted gene protein sequences were aligned using MAFFT, using the local pair option (mafft-linsi), the tree was calculated using iqTree (Nguyen et al., 2015). The unrooted trees (**Figs. S8-13, S15**) were visualized using the online web tool from the Interactive Tree of Life (IToL, https://itol.embl.de) and subsequently manually colored. The rooted tree of the full cdh gene cluster (**Fig. S14**) was calculated with iqtree, using the concatenated sequence of separately aligned subunits (mafft-linsi). The cdh gene cluster of OP bin 004 (a Deltaproteobacterium) was used as the outgroup.

## Additional information

### Acknowledgements

The authors would like to acknowledge members of the Quake Lab Sequencing Facility including Jennifer Okamoto and Norma Neff as well as the Department of Energy (DOE) Joint Genome Institute’s (JGI) assembly and annotation teams. We would also like to thank Anastasia Nedderton (Stanford) and NPS staff at YNP including Research Coordinator Christie Hendrix for assistance with different aspects of the sample collection, preservation, and characterization processes. This work is supported by the Templeton Foundation. The work conducted by the U.S. Department of Energy Joint Genome Institute, a DOE Office of Science User Facility, is supported under Contract No. DE-AC02-05CH11231. BAB is supported by a Rubicon Fellowship from the Netherlands Organization for Scientific Research (NWO), and PCB is supported by the Burroughs Welcome Fund via a Career Award at the Scientific Interface.

### Author contributions

Sample collection: PCB. Conceptualization: SRQ, PCB, FBY, BAB. Performing (mini-)metagenomic experiments, library preparation and sequence assembly: BAB, FBY. Binning and quality control: BAB, FBY. Metabolic reconstruction and analysis: BAB. Data interpretation: BAB, FS, TW, SRQ. Genome tree placement and interpretation: FS, BAB, TW. Writing the manuscript (original draft): BAB. Reviewing and editing the manuscript: BAB, FS, PCB, TW, SRQ.

### Author ORCIDs

Bojk A. Berghuis, https://orcid.org/0000-0002-7797-0553

Feiqiao Brian Yu, http://orcid.org/0000-0003-3416-3046

Frederik Schulz, http://orcid.org/0000-0002-4932-4677

Paul C. Blainey, https://orcid.org/0000-0002-4889-8783

Tanja Woyke, http://orcid.org/0000-0002-9485-5637

Stephen R. Quake, http://orcid.org/0000-0002-1613-0809

